# Ran GTPase regulates non-centrosomal microtubule nucleation and is transported by actin waves towards the neurite tip

**DOI:** 10.1101/684720

**Authors:** Yung-An Huang, Chih-Hsuan Hsu, Ho-Chieh Chiu, Chris T. Ho, Wei-Lun Lo, Eric Hwang

**Affiliations:** Department of Biological Science and Technology, National Chiao Tung University, Hsinchu, Taiwan; Institute of Molecular Medicine and Bioengineering, National Chiao Tung University, Hsinchu, Taiwan; Institute of Bioinformatics and Systems Biology, National Chiao Tung University, Hsinchu, Taiwan; Center for Intelligent Drug Systems and Smart Bio-devices (IDS2B), National Chiao Tung University, Hsinchu, Taiwan

## Abstract

Microtubule (MT) is the most abundant cytoskeleton in neurons and controls multiple facets of their development. While the organizing center of MTs in mitotic cells is typically located at the centrosome, MT nucleation in post-mitotic neurons switches to non-centrosomal sites. A handful of proteins and organelle have been shown to promote non-centrosomal MT formation in neurons, yet the regulation mechanism remains unknown. Here we demonstrate that the small GTPase Ran is a key regulator of non-centrosomal MT nucleation in neurons. The GTP-bound Ran (RanGTP) localizes to the neurite tips and around the soma. Using the RanGTP- and RanGDP-mimic mutants, we show that RanGTP promotes MT nucleation at the tip of the neurite. To demonstrate that RanGTP can promote MT nucleation in regions other than the neurite tip, an optogenetic tool called RanTRAP was constructed to enable light-induced local production of RanGTP in the neuronal cytoplasm. An increase of non-centrosomal MT nucleation can be observed by elevating the RanGTP level along the neurite using RanTRAP, establishing a new role for Ran in regulating neuronal MTs. Additionally, the mechanism of RanGTP enrichment at the neurite tip was examined. We discovered that actin waves drive the anterograde transport of RanGTP towards the neurite tip. Pharmacological disruption of actin waves abolishes the enrichment of RanGTP and reduces the non-centrosomal MT nucleation at the neurite tip. These observations provide a novel regulation mechanism of MTs and an indirect connection between the actin and MT cytoskeletons in neurons.

## Introduction

The nervous system is the main information relaying and processing unit in multicellular organisms to interact with the external world. In order for the nervous system to operate, individual neurons must be connected in a highly ordered manner. To achieve such organized connections, the developmental processes following the generation of a terminally differentiated neuron must be highly regulated. It has been shown that most if not all developmental processes of neurons depend on the organization and function of an essential cellular fibrous network called microtubule cytoskeleton. MTs are tube-like polymers composed of heterodimers of α- and β-tubulins, they are also highly dynamic polymers that utilize GTP hydrolysis to control polymerization and depolymerization as well as the transition between the two phases (Brouhard and Rice, 2018; Desai and Mitchison, 1997). The ordered assembly of α/β-tubulin heterodimers gives MTs two distinct ends: a dynamic plus-end where polymerization and depolymerization occur, and an inert minus-end where nucleation event happens. In cells actively undergoing proliferation, MT nucleation usually takes place at the centrosomes. Differentiated cells, on the other hand, contain largely of non-centrosomal MTs (ncMTs) that are not organized at the centrosome (Bartolini and Gundersen, 2006; Keating and Borisy, 1999; Muroyama and Lechler, 2017; Sanchez and Feldman, 2017). In neurons, MTs are initially assembled from the centrosome (Yu et al., 1993). But as neurons mature, the centrosome loses its MT-organizing center (MTOC) capability (Leask et al., 1997; Stiess et al., 2010). Several cellular components have been identified as the non-centrosomal MT-organizing center (ncMTOC) in neurons. Golgi outposts was the first to be demonstrated for nucleating ncMTs in dendrites of *Drosophila* da neurons (Ori-McKenney et al., 2012). Although it is important to point out that Golgi outposts are not present in all dendrites and genetically forcing Golgi outposts out of dendrites does not affect the MT organization (Nguyen et al., 2014). The augmin complex was later shown to nucleate ncMTs from the existing MTs in both the axonal and dendritic compartments (Cunha-Ferreira et al., 2018; Sanchez-Huertas et al., 2016). In line with the augmin complex discovery, another protein (TPX2) required for branch MT formation on existing MTs (Petry et al., 2013) has also been shown to promote ncMT nucleation in both axons and dendrites of mammalian neurons (Chen et al., 2017). Interestingly, the small GTPase Ran which plays an important role in regulating TPX2 activity during mitosis has been reported to regulate TPX2-mediated ncMT nucleation in neurons (Chen et al., 2017).

The Ras-related nuclear protein (Ran) is a member of the Ras superfamily GTPase that is crucial in the process of nucleocytoplasmic transport (Gorlich and Mattaj, 1996). In addition to its role in nucleocytoplasmic transport, Ran has also been shown to affect spindle formation in *Xenopus laevis* egg extract (Kahana and Cleveland, 1999). The effect of Ran on mitotic spindle formation is mediated by importin-α/β heterodimer, which binds to nuclear localization sequence (NLS) on spindle assembly factors (SAFs) and inhibits their activity (Clarke and Zhang, 2008). In the presence of RanGTP, SAFs are released from the inhibitory importin heterodimer and allowed to promote MT nucleation to facilitate the assembly of the mitotic spindle. One of the SAFs is the aforementioned TPX2 (Gruss et al., 2001), which can promote branching MT nucleation from existing MTs (Petry et al., 2013). Although the effect of Ran on MT nucleation is well established, most of the studies were carried out in meiotic egg extract or mitotic cells. The effect of Ran on MTs of post-mitotic neurons has received much less attention. A few lines of evidence indicate that Ran plays a role in neuronal morphogenesis. First, Ran depletion in primary *Drosophila* neurons displayed excessive neurite branching and blebbing (Sepp et al., 2008). Second, Ran or RCC1 (the Ran guanine nucleotide exchange factor) knockdown compromises axon specification in mammalian neurons (Mencarelli et al., 2018). Third, RanGTP hydrolysis in the axoplasm has been observed after the nerve injury and perturbing the hydrolysis of RanGTP compromises the regeneration of axons (Yudin et al., 2008). These data indicate that Ran is an important regulator of neuronal morphogenesis under both normal and injured conditions.

A recent discovery shows that RanGTP is specially enriched at the tip of the neurite and around the soma (Chen et al., 2017). When a pharmacological perturbation disrupts the interaction between Ran and its downstream effector, ncMT nucleation from the neurite tip is compromised. These observations raise the questions whether Ran GTPase can regulate ncMT formation from anywhere along the neurite and what is the mechanism of RanGTP enrichment at the neurite tip. To answer these questions, we constructed an optogenetic reagent called RanTRAP which enables the local increase of RanGTP at the photoactivation site. By photoactivating RanTRAP at specific region along the neurite, we show that RanGTP can indeed promote ncMT formation. In addition, we detected a colocalization of RanGTP and actin wave in neurons. By examining the motility of a RanGTP-mimic mutant, we discovered that RanGTP is transported anterogradely by actin waves towards the neurite tip. Pharmacological disruption of actin waves reduces the level of RanGTP as well as decreases the emanating frequency of MTs at the neurite tip. These observations confirm the role of Ran GTPase in regulating ncMT nucleation and provide a novel mechanism of moving the active RanGTP molecules towards the neurite tip where ncMT nucleation occurs.

## Results

### Ran GTPase affects microtubule formation at the neurite tip

It has been reported that RanGTP is enriched at the soma and the neurite tip (Chen et al., 2017). To confirm this localization, an antibody which specifically targets the C-terminal tail of Ran GTPase that is only exposed in the GTP-bound state (Richards et al., 1995) was used for immunofluorescence staining to examine 2 days *in vitro* (2DIV) dissociated hippocampal neurons. Similar to the previous observation, RanGTP localizes to the tips of both axon and dendrite and around the soma (Figure 1A). In addition, the tip of the dendrite exhibits a higher level of RanGTP than that of the axon (Figure 1B). Interestingly, RanGTP appears to colocalize with the actin-based structure in the growth cone (Figure 1C).

**Figure 1.**
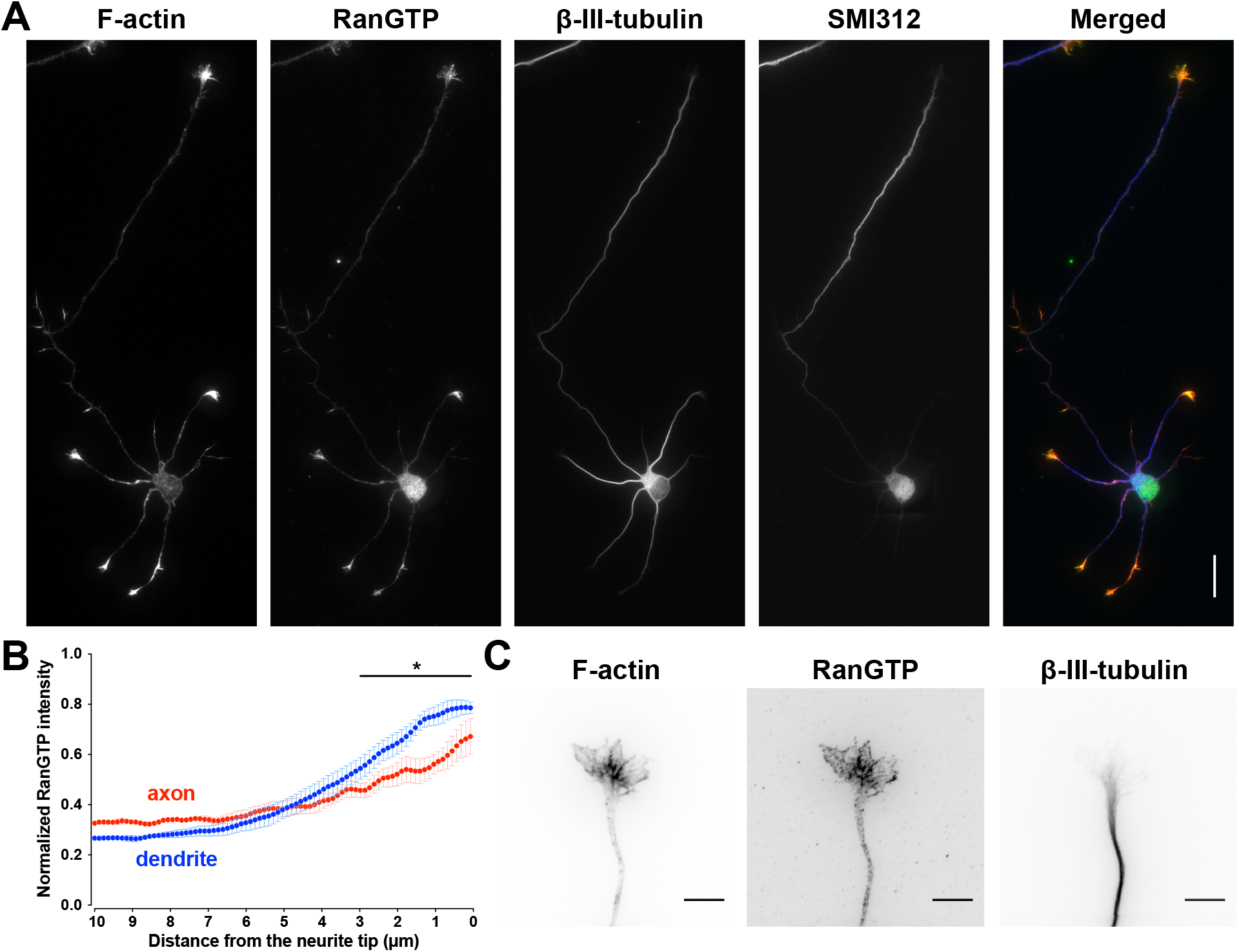
GTP-bound Ran is enriched at both axon and dendrite tips, and colocalized with actin-based structures. (A) Representative images of a 2DIV hippocampal neuron immunofluorescence stained with RanGTP, β-III-tubulin, SMI312 antibodies, and phalloidin. The merged image shows phalloidin staining in red, RanGTP in green, and β-III-tubulin in blue. The scale bar represents 25 μm. (B) RanGTP intensity linescan along a 10 μm stretch from axon (red) or dendrite (blue) tips in 2DIV hippocampal neurons. Error bars represent SEM from 3 independent experiments, * *p*<0.05, two-way ANOVA followed by Bonferroni post-hoc test. (C) Representative image of the growth cone from a 2DIV hippocampal neuron fixed and stained with RanGTP (middle), β-III-tubulin (right) antibody, and phalloidin (left). Images were inverted to facilitate visualization. Scale bars represent 10 μm.

Cytoplasmic RanGTP has also been shown to activate TPX2 and to promote ncMT nucleation in neurons (Chen et al., 2017). To examine whether RanGTP regulates ncMT nucleation, Ran mutants were utilized to alter the level of cytoplasmic RanGTP in neurons. The constitutively active Ran mutant (RanQ69L) which mimics RanGTP is used to increase cytoplasmic RanGTP, while the dominant negative Ran mutant (RanT24N) which irreversibly binds to the Ran guanine nucleotide exchange factors (RanGEF) is used to reduce cytoplasmic RanGTP (Klebe et al., 1995). We first examined whether expressing RanQ69L or RanT24N can alter the level of cytoplasmic RanGTP in neurons. Interestingly, expressing RanQ69L or RanT24N specifically alters RanGTP level at the neurite tip (Figure S1) without affecting its level along the entire neurite length (data not shown). We then examined MT formation at the neurite tip in neurons expressing these Ran mutants. The CNS-enriched MT plus-end tracking protein EB3 was used to assess the formation of MTs (Nakagawa et al., 2000). As expected, neurons expressing RanQ69L exhibit a significant increase of MT formation frequency at their neurite tips compared to those expressing wild-type Ran or RanT24N; neurons expressing RanT24N exhibit a significant decrease of MT formation frequency compared to wild-type Ran or RanQ69L-expressing neurons (Figure 2). On the other hand, the time MT stays in the polymerization phase (persistence) and the MT polymerization rate are not affected. In addition, we discovered that Ran mutants expression can alter neuronal morphogenesis (Figure S2). It is of interest to point out that neurons expressing AcGFP-fused wild-type Ran show decreased neurite length compared to those expressing cytosolic AcGFP, suggesting an overabundance of wild-type Ran can negatively affect neurite elongation. Neurons expressing the RanGTP-mimic mutant (RanQ69L) extend longer neurites than those expressing wild-type Ran; while neurons expressing RanGDP-mimic mutant (RanT24N) possess shorter neurites than wild-type Ran-expressing ones. Neurite branching is not significantly altered by the expression of Ran mutants, this is probably due to the low branching capability at such an early stage. On the other hand, neurons expressing RanT24N sprout fewer primary neurites than those expressing RanQ69L or wild-type Ran. Taken together, these data indicate changing the level of RanGTP at the neurite tip alters the ncMT nucleation at this location and affects the overall morphology of the neuron.

**Figure 2.**
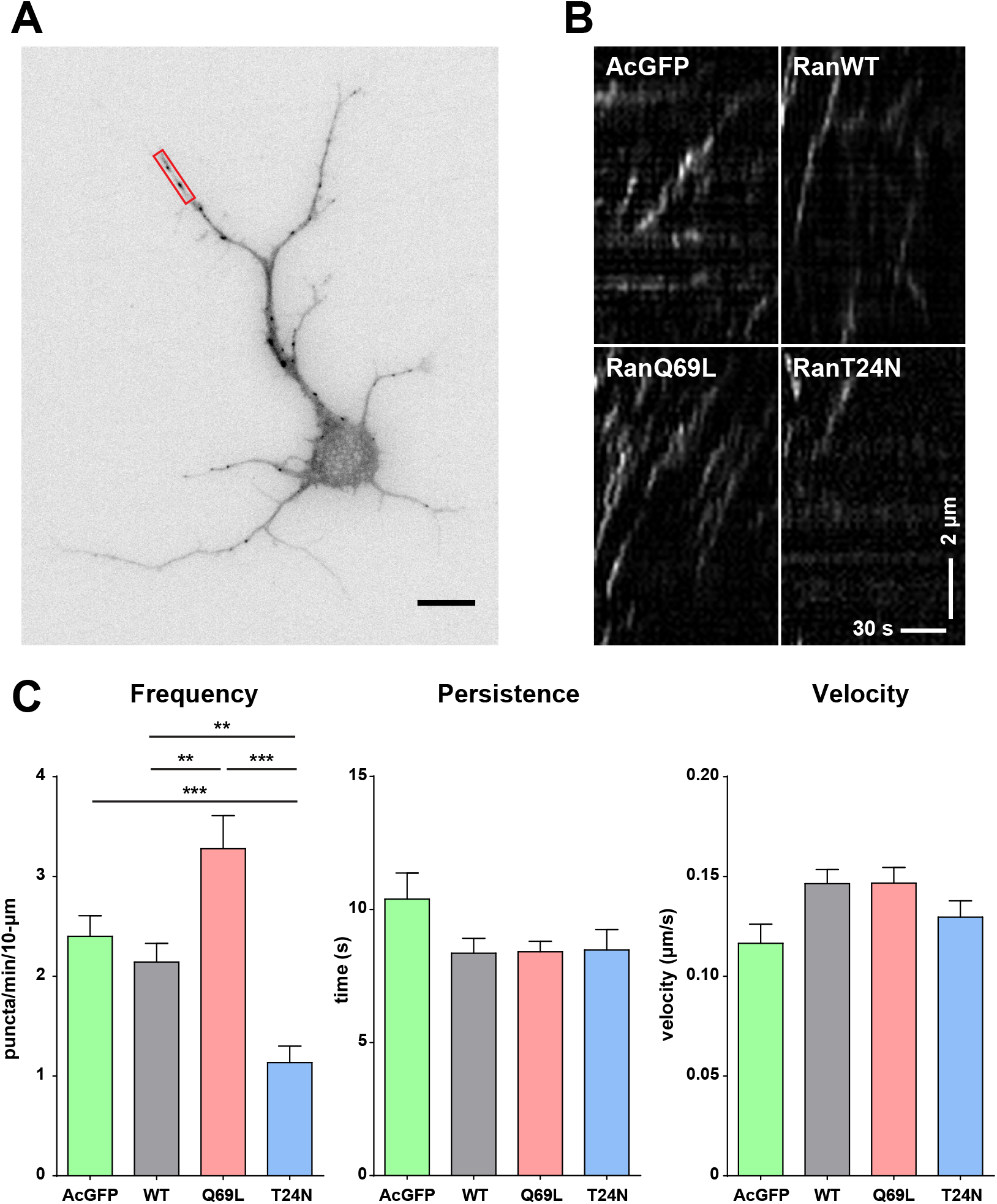
Ran mutants affect microtubule nucleation at the neurite tip. (A) Representative images of 2DIV dissociated cortical neurons co-expressing AcGFP-RanWT and EB3-mCherry; only the EB3-mCherry channel is shown. The red window indicated the region where kymographs were generated. The scale bar represents 10 μm. (B) Representative kymographs of EB3-mCherry at the tip of the neurite in various Ran mutants expressing neurons. (C) Quantification of EB3-mCherry dynamics in AcGFP (green), AcGFP-RanWT (gray), AcGFP-RanQ69L (red), and AcGFP-RanT24N (blue) expressing neurons. More than 8 neurites, 6 neurons were analyzed per condition. ** *p*<0.01, *** *p*<0.001, one-way ANOVA followed by Tukey’s post-hoc test. Error bars represent SEM from 3 independent experiments.

### A photoactivatable Ran promotes microtubule formation along the neurite

We have shown that expressing Ran mutants alters the level of cytoplasmic RanGTP at the neurite tip and leads to the change of ncMT nucleation at this location. To demonstrate that Ran GTPase is a major regulator of ncMT nucleation, it will be important to show that RanGTP can induce ncMT formation at other regions of the neuron. Since altering the level of cytoplasmic RanGTP can lead to detrimental effect on cell survival due to its influence on nucleocytoplasmic transport, it is therefore crucial to have the capability of controlling the location of RanGTP production with spatial precision and examine its effect on ncMT nucleation. To achieve this goal, we utilized the LOVTRAP system developed in Klaus Hahn’s lab (Wang et al., 2016). This system takes advantage of a photoreactive domain called LOV2 that derived from phototropins in plants (Huala et al., 1997; Salomon et al., 2000). In the absence of light, LOV2 domain binds to a short peptide derived from the Z subunit of protein A called Zdark (ZDK). Upon blue light irradiation, a conformational change in the LOV2 domain causes it to dissociate from ZDK. By fusing LOV2 domain to the organelle targeting sequence and fusing ZDK to the protein of interest, one can sequester this protein of interest to a specific organelle in the dark and release it at specific time and location upon light irradiation. We used this system to design an optogenetic tool called “RanTRAP” to spatially control the level of RanGTP. Our RanTRAP system is composed of two parts: a mitochondrial targeting sequence fused to the LOV2 domain (NTOM20-LOV2) and the aforementioned RanGTP-mimic RanQ69L fused to the ZDK (mCherry-ZDK-RanQ69L). In the absence of light, mCherry-ZDK-RanQ69L binds to NTOM20-LOV2 and sequesters the activity of this RanGTP-mimic mutant to the mitochondria. Upon light irradiation, LOV2 and ZDK dissociate from each other, releasing mCherry-ZDK-RanQ69L from the mitochondria. mCherry-ZDK-RanQ69L then binds to importin-β and disrupts the inhibitory importin complex. This leads to the activation of TPX2 and the nucleation of ncMT (Figure 3A). In addition to the RanGTP-mimic RanQ69L, the RanGDP-mimic RanT24N was also fused to ZDK to serve as the control. These constructs are referred to as RanQ69L-TRAP and RanT24N-TRAP hereafter.

**Figure 3.**
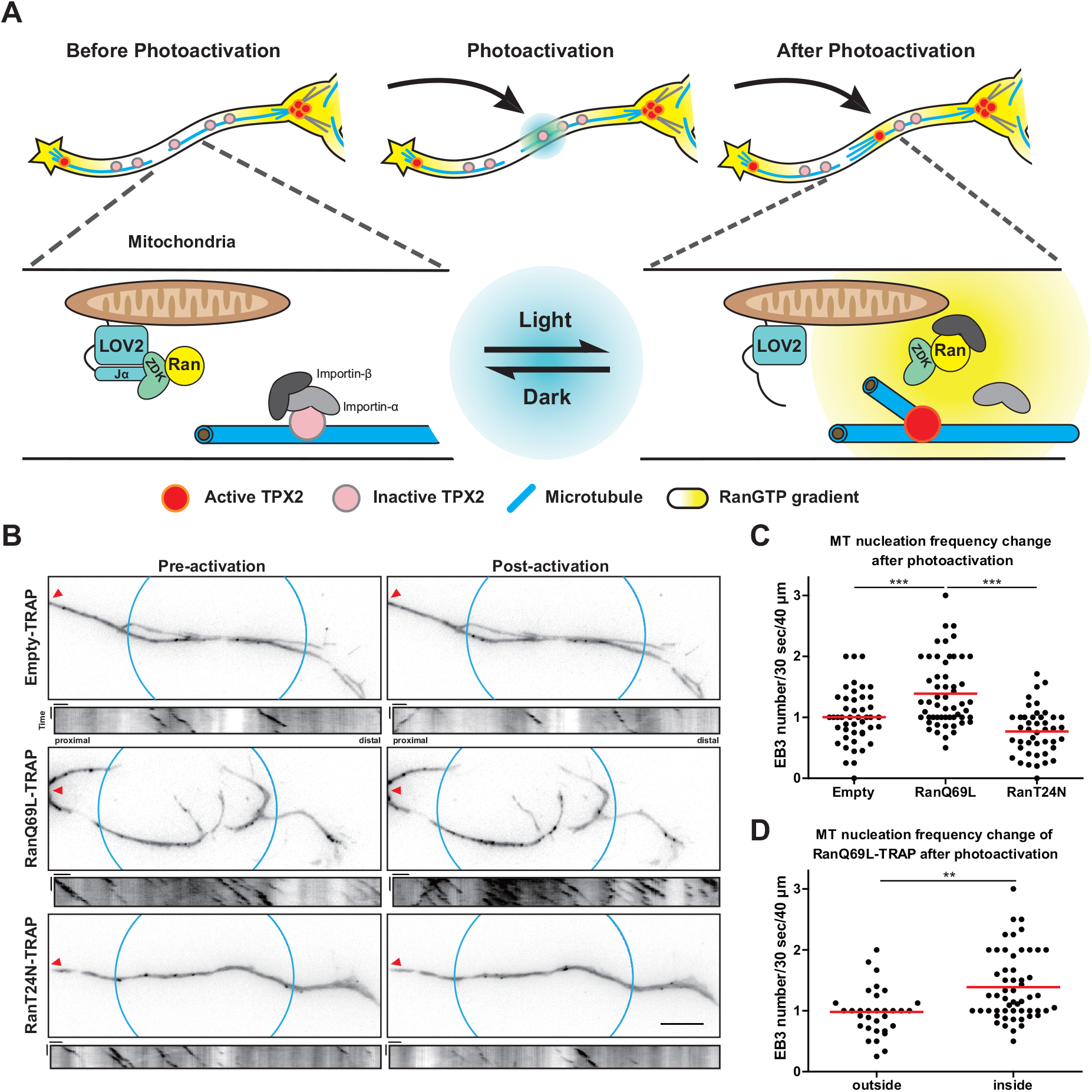
Locally producing RanGTP promotes non-centrosomal microtubule nucleation along the neurite. (A) Schematic illustration of locally photoactivating RanTRAP in promoting non-centrosomal microtubule nucleation along the neurite. (B) Representative images and kymographs of 4DIV mouse hippocampal neurons expressing EB3-mCherry and Empty-TRAP (top), RanQ69L-TRAP (middle), or RanT24N-TRAP (bottom). Blue circles indicate the region of photoactivation and kymographs generation. The red arrowheads denote the proximal side of the neurite. All images have the same scale and the scale bar represents 10 μm. In all kymographs, vertical scale bars represent 15 seconds and horizontal scale bars represent 2 μm. (C) Quantification of EB3-mCherry frequency in photoactivated regions after photoactivation. The red horizontal bars indicate the mean frequency. This result was obtained from three or four (4 for RanQ69L-TRAP group) independent repeats. *** *p*<0.001, one-way ANOVA followed by Tukey’s post-hoc test. (D) Quantification of EB3-mCherry frequency inside or outside the photoactivation region in neurons expressing RanQ69L-TRAP and EB3-mCherry. The red horizontal bars indicate the mean frequency. This result was obtained from four independent experiments. ** *p*<0.01, two-tailed Student’s *t*-test.

We first validated this RanTRAP platform by examining whether it can mask the phenotypic effect of RanQ69L or RanT24N expression in mitotic cells. It has been shown that overexpressing the constitutively active RanQ69L or the dominant negative RanT24N leads to the formation of abnormal mitotic spindles (Moore et al., 2002). We reason that if this RanTRAP tool can sequester Ran mutants on mitochondria, these mutants would not be able to cause the formation of abnormal spindle. To increase the mitotic index of HeLa cells, double thymidine arrest was utilized to synchronize these cells (Figure S3A). The mitotic spindle morphology and chromosome alignment in mitotic HeLa cells expressing RanQ69L-TRAP or RanT24N-TRAP were then examined. Abnormal mitotic spindles were classified into three main categories: 1) bipolar spindle with misaligned chromosomes, 2) multipolar spindle, or 3) monopolar spindle (Figure S3B). In cells expressing mCherry-ZDK-RanQ69L or mCherry-ZDK-RanT24N alone, the percentage of HeLa cells with abnormal spindle is significantly higher than untransfected cells, demonstrating that mCherry-ZDK-Ran mutants cause abnormal mitotic spindle formation in our hands. In contrast, cells expressing NTOM20-LOV2-WT (NTOM20 fused to the wild-type LOV2) exhibit low percentage of abnormal spindle similar to untransfected cells. In cells co-expressing NTOM20-LOV2-WT and mCherry-ZDK-Ran mutant, the percentage of cells with abnormal spindle is significantly lower than those expressing mCherry-ZDK-Ran mutant alone or those co-expressing NTOM20-LOV2-I539E (a mutant LOV2 that always stays in the photoactivated conformation) (Harper et al., 2004) and mCherry-ZDK-Ran mutant (Figure S3C), indicating that sequestering Ran protein onto the mitochondria can block its function. These results demonstrate that RanTRAP does inhibit the function of Ran mutants in the absence of light.

We then examined whether RanTRAP can target Ran mutants onto mitochondria along the neurite. When the plasmid expressing NTOM20-mVenus-LOV2-WT was introduced into mouse cortical neurons, it colocalized with MitoTracker signal in the neurite (Figure S4A), indicating that NTOM20-mVenus-LOV2-WT can be readily targeted to the mitochondria. When the plasmid expressing NTOM20-mVenus-LOV2-WT and the plasmid expressing mCherry-ZDK-Ran were co-transfected into dissociated neurons, colocalization can also be detected along the neurite (Figure S4B). These results demonstrate that mitochondrion is an ideal organelle to target and sequester Ran mutant along the neurite. Furthermore, it has been documented that the molar ratio of LOV2- to ZDK-fused proteins is crucial for achieving maximal release of the trapped protein upon irradiation (Wang and Hahn, 2016). Since the molar ratio of LOV2- and ZDK-fused proteins cannot be readily determined, the molar ratio their expression plasmids was empirically tested to achieve maximal release upon irradiation (Figure S5).

Next, we sought to determine whether locally photoactivate RanTRAP-expressing neurons can release RanGTP-mimic mutant in light irradiated region of the neurite. In this experiment, plasmids expressing NTOM20-mVenus-LOV2-WT and mCherry-ZDK-RanQ69L (RanQ69L-TRAP) were transfected into dissociated neurons. The dynamic localization of mCherry-ZDK-RanQ69L before and after a local photoactivation was then examined. Upon local photoactivation, mCherry-ZDK-RanQ69L was immediately released from the mitochondria into the cytosol. This cytosolic pool of mCherry-ZDK-RanQ69L then gradually reassociated back to the mitochondria (Figure S6). This result shows that spatial and reversible release of RanGTP along the neurite can be achieved in dissociated neurons.

Finally, to confirm whether the cytoplasmic RanGTP can promote ncMT nucleation in neurons, the photoactivatable RanTRAP system was utilized to locally release RanGTP at the photoactivated region along the neurite. We reason that if RanGTP does promote ncMT nucleation, releasing the RanGTP-mimic RanQ69L will cause an increase of MT formation at the photoactivation site (Figure 3A). To examine MT nucleation, EB3-mCherry was used to label the growing MT plus-ends. A 24-pulse regime (24 pulses of 80 milliseconds irradiation with 5 seconds duration in between pulses) was established to generate a sustained level of RanGTP for ~2 minutes at the photoactivation region. This 2-minute time period was selected based on the observation that adding RanQ69L and TPX2 dramatically enhances MT nucleation within 66 seconds in *Xenopus* egg extract (Petry et al., 2013). To maximize the effect of RanTRAP, photoactivation was carried out in the region of the neurite more than 15 µm from the soma or the neurite tip, where endogenous RanGTP level is low (Chen et al., 2017). EB3-mCherry dynamics was quantified before and after photoactivation (Figure 3B). As expected, photoactivating neurons co-expressing NTOM20-mVenus-LOV2-WT and mCherry-ZDK-RanQ69L (RanQ69L-TRAP) leads to a significant increase of EB3-mCherry comets in the irradiated region compared to photoactivating neurons co-expressing NTOM20-mVenus-LOV2-WT and mCherry-ZDK (Empty-TRAP) (Figure 3C). Furthermore, the increase of EB3-mCherry comets after photoactivation only occurs in the irradiated region (Figure 3D). On the other hand, photoactivating neurons expressing NTOM20-mVenus-LOV2-WT and the RanGDP-mimic mCherry-ZDK-RanT24N (RanT24N-TRAP) does not significantly alter the frequency of EB3-mCherry comets (Figure 3C). These results demonstrate that locally releasing RanGTP by itself can enhance the nucleation of ncMT along the neurite, suggesting that the cytoplasmic RanGTP is the major regulator of ncMT in neurons.

### Actin waves transport GTP-bound Ran towards the neurite tip

The localization of RanGTP at the neurite tip (Figure 1) prompts us to determine the mechanism of peculiar tip enrichment. Two possible mechanisms can explain the tip enriched RanGTP localization. In the first mechanism, an unknown RanGEF localizes to the neurite tip and locally produces the RanGTP. In the second mechanism, the RanGTP molecules are produced elsewhere (potentially in the soma) and transported towards the neurite tip. Our current data argue against the first mechanism, because a RanGTP-mimic mutant is enriched while a RanGDP-mimic mutant is essentially absent at the neurite tip when expressed in neurons (Figure S1). Since RanGEF has a higher affinity towards RanGDP, a tip localized RanGEF should cause the enrichment of RanGDP instead of RanGTP at the neurite tip. On the other hand, several observations support the RanGTP transportation mechanism. First, we detected the colocalization of RanGTP with actin-based structure in the growth cone (Figure 1C). Second, MT nucleation frequency has been observed to increase immediately behind actin waves (Winans et al., 2016). Given that actin waves are responsible for the anterograde transport of a variety of biological molecules (Inagaki and Katsuno, 2017), we hypothesize that they are also in charge of moving RanGTP molecules towards neurite tips. Perhaps the elevated MT nucleation frequency observed by Winan and colleagues is caused by RanGTP moving with actin waves, causing ncMT formation in its wake. As a first step towards confirming this RanGTP transportation mechanism, we examined the involvement of the actin cytoskeleton on the tip localization of RanGTP. 1DIV dissociated neurons were treated with 2.5 μM cytochalasin D for 6 hours to depolymerize actin filaments, this treatment condition has been shown to compromise actin-based structures in the neurite without affecting neurite formation or elongation (Chia et al., 2016). While actin filaments in DMSO-treated control neurons are unaffected, they are largely absent from the growth cone in cytochalasin D-treated neurons (Figure 4A and B). In addition, RanGTP is no longer enriched at the neurite tip in cytochalasin D-treated neurons (Figure 4C and D). This result indicates that the actin-based structures are responsible for the enrichment of RanGTP at the neurite tip.

**Figure 4.**
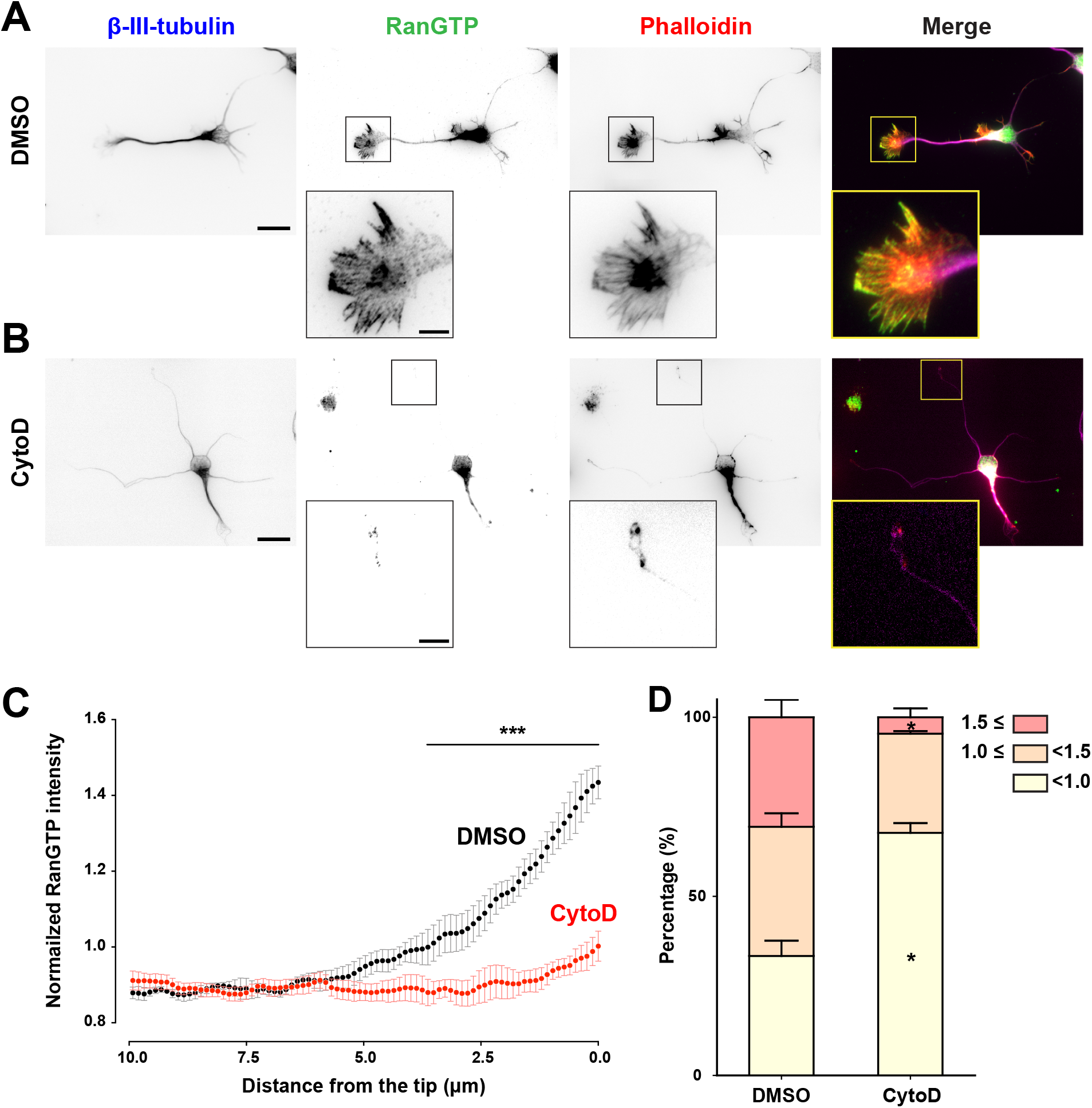
The localization of RanGTP at the neurite tip depends on the actin cytoskeleton. Representative images of 1DIV hippocampal neurons treated with DMSO (A) or 2.5 μM cytochalasin D (B) for 6 hours. The boxed areas at the neurite tips are magnified in the insets. The scale bars represent 20 μm and 5 μm in insets. (C) Normalized RanGTP level linescans along a 10 μm stretch from the neurite tip in DMSO (black) and 2.5 μM cytochalasin D (red) treated neurons. RanGTP intensity is normalized to the mean intensity along the entire neurite. The dots and error bars indicate mean and SEM from 3 independent experiments. *** *p*<0.001, two-way ANOVA followed by Bonferroni post-test. (D) Quantification of RanGTP intensity ratio at the neurite tip. The mean intensity of RanGTP within 1 μm from the neurite tip was divided by the intensity along the entire neurite. RanGTP intensity ratios are classified into three groups: the group with ratio equal to or greater than 1.5 is shown in red, the group with ratio between 1.5 and 1.0 is shown in orange, and the group with ratio below 1.0 is shown in yellow. * *p*<0.05, two-tailed Student’s *t*-test. Error bars represent SEM from three independent experiments, with a total of 135 and 141 neurites analyzed in the DMSO or cytochalasin D treated neurons, respectively.

Given that the actin wave has been documented to transport various cellular cargoes in neurons (Inagaki and Katsuno, 2017) and that neurite tip enriched RanGTP depends on actin-based structures, the actin wave appears as an attractive candidate for the localization of RanGTP. To determine whether actin waves are the driving force for the anterograde transport of RanGTP, we first examined the localization of RanGTP and actin waves in fixed neurons. In addition to its high abundance in the growth cone, RanGTP can also be detected to colocalize with the actin wave (Figure 5A and 5B). Consistent with the previous observation (Winans et al., 2016), β-III-tubulin signal intensity can be seen to increase in and behind the actin wave (Figure 5B). In addition to fixed neurons, we examined the motility of RanQ69L (RanGTP-mimic) in young neurons that are actively generating actin waves (Flynn et al., 2009; Ruthel and Banker, 1999). Plasmid expressing AcGFP-RanQ69L, AcGFP-RanT24N, or cytosolic AcGFP was introduced into dissociated neurons just before plating and the motility of the AcGFP signal as well as the actin waves was monitored at 2DIV using live cell imaging. Consistent with our hypothesis, AcGFP-RanQ69L can be observed to move anterogradely in clusters along the neurite and co-migrate with the actin wave towards the tip (Figure 5C and 5F). In contrast, AcGFP-RanT24N exhibits minimal signal in the neuronal cytoplasm and does not co-migrate with the actin wave (Figure 5D and 5G). To eliminate the possibility that the co-migration of RanQ69L and actin wave is due to the increase of cytoplasmic volume around the wave, we also examined the motility of cytosolic AcGFP (Figure 5E and 5H). While the cytosolic AcGFP appears to migrate with the actin wave, its movement is less persistent and its enrichment in the wave is also less pronounced than that of AcGFP-RanQ69L. To quantify the co-migration between Ran mutant and the actin wave, we measured the AcGFP signal in a selected region of interest (ROI) along the neurite that experienced an actin wave from 15 minutes before to 15 minutes after the arrival of the actin wave. The time at which the actin wave passes through the ROI is set to 0 minute (Figure 5I). If the AcGFP-tagged protein co-migrates with the actin wave, one would expect the signal intensity of AcGFP to peak at the time of actin wave passage. This is indeed the case for AcGFP-RanQ69L (Figure 5J). In contrast, AcGFP-RanT24N signal does not peak at 0 minute. It should be noted that the large fluctuation of AcGFP-RanT24N signal over time is due to its low intensity in the neuronal cytoplasm. Interestingly, while cytosolic AcGFP exhibits peak intensity when the actin wave passes through the ROI, its relative intensity at the peak is significantly lower than that of AcGFP-RanQ69L. This indicates that AcGFP-RanQ69L is more enriched in the actin wave than the cytosolic AcGFP molecule. Taken together, our data demonstrate that RanGTP is transported by actin waves towards the neurite tip.

**Figure 5.**
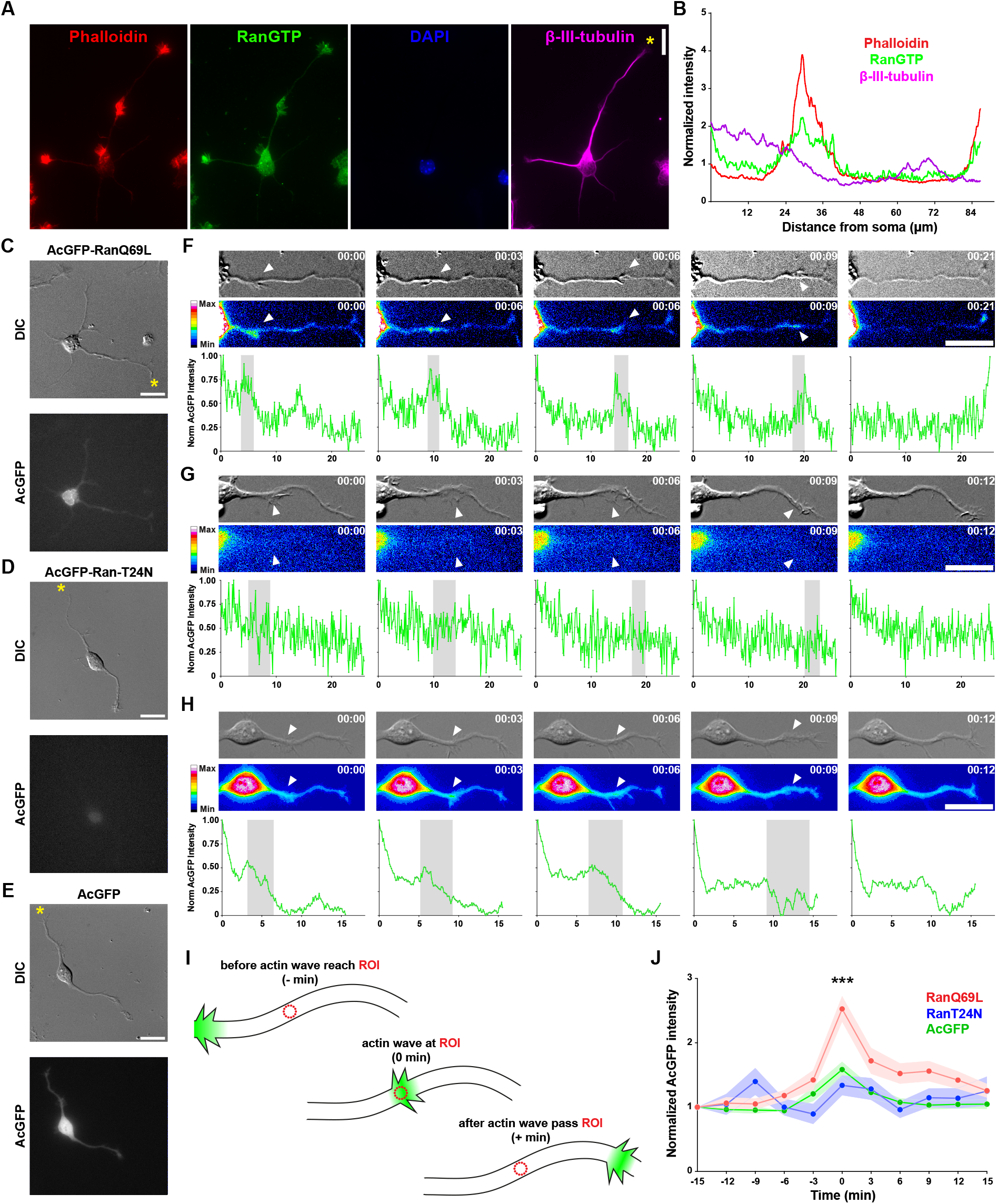
GTP-bound Ran is transported by the actin wave towards the neurite tip. (A) Representative images of 1DIV hippocampal neurons immunofluorescence stained with antibodies against RanGTP (green) and the neuron-specific β-III-tubulin (purple). Actin filaments and the nucleus were stained with phalloidin (red) and DAPI (blue), respectively. The scale bar represents 20 μm. (B) The intensity linescan of indicated molecules along the asterisked neurite in (A). (C-E) Representative images of actin waves-containing 2DIV cortical neurons expressing AcGFP-RanQ69L (C), AcGFP-RanT24N (D), AcGFP (E). The scale bars represent 10 μm. (F-H) Time-lapse DIC (top) and AcGFP (middle) images, as well as the AcGFP intensity linescan (bottom) of a single neurite in the neuron expressing AcGFP-RanQ69L (F), AcGFP-RanT24N (G), or AcGFP (H). Neurite segments in (F), (G), (H) are derived from the asterisked neurite in (C), (D), (E), respectively. The white arrowheads mark the location of the actin wave. The time stamps (hour:min) indicate the time progressed since the first image. The gray shaded area in the linescan graphs indicates the location of the actin wave. The scale bars represent 10 μm. (I) Schematic illustration of the method for quantifying the AcGFP intensity fluctuation over time in a selected ROI along the neurite shaft. The dotted circles indicate the selected ROI where AcGFP signal (green glow) was measured from. (J) Quantification of AcGFP intensity fluctuation in the selected ROI along the neurite shaft from 15 minutes before to 15 minutes after the arrival of the actin wave. The time at which the actin wave passes through the selected ROI is set to 0 minute. *** *p*< 0.001 two-way ANOVA followed by Tukey post-hoc analysis (between RanQ69L and RanT24N as well as between RanQ69L and AcGFP). Dots and shaded areas indicate mean and SEM. More than 22 actin wave-containing neurites were analyzed in each condition from three independent repeats.

### Disrupting actin waves reduces microtubule formation at the neurite tip

Given that RanGTP is responsible for promoting ncMT nucleation and it is transported by actin waves towards the neurite tip, the logical outcome of disrupting the propagation of actin wave is the reduction of MT nucleation at the tip of the neurite. We set out to test this hypothesis by treating neurons with 2.5 μM cytochalasin D for 6 hours, a condition previously shown to eliminate actin waves and significantly reduce RanGTP at the tip (Figure 4). Differential interference contrast (DIC) microscopy was used to identify the tip of the neurite. While cytochalasin D treatment does not affect MT nucleation around the soma (Figure 6A), it causes a significant reduction of MT nucleation at the neurite tip (Figure 6B and 6C). Furthermore, cytochalasin D treatment only reduces the frequency of MT nucleation while leaving MT polymerization velocity or the time MT spent in the polymerization phase unaffected (Figure 6C). This result shows that ncMT nucleation at the neurite tip is influenced by the actin wave, providing a new indirect connection between these two cytoskeletons.

**Figure 6.**
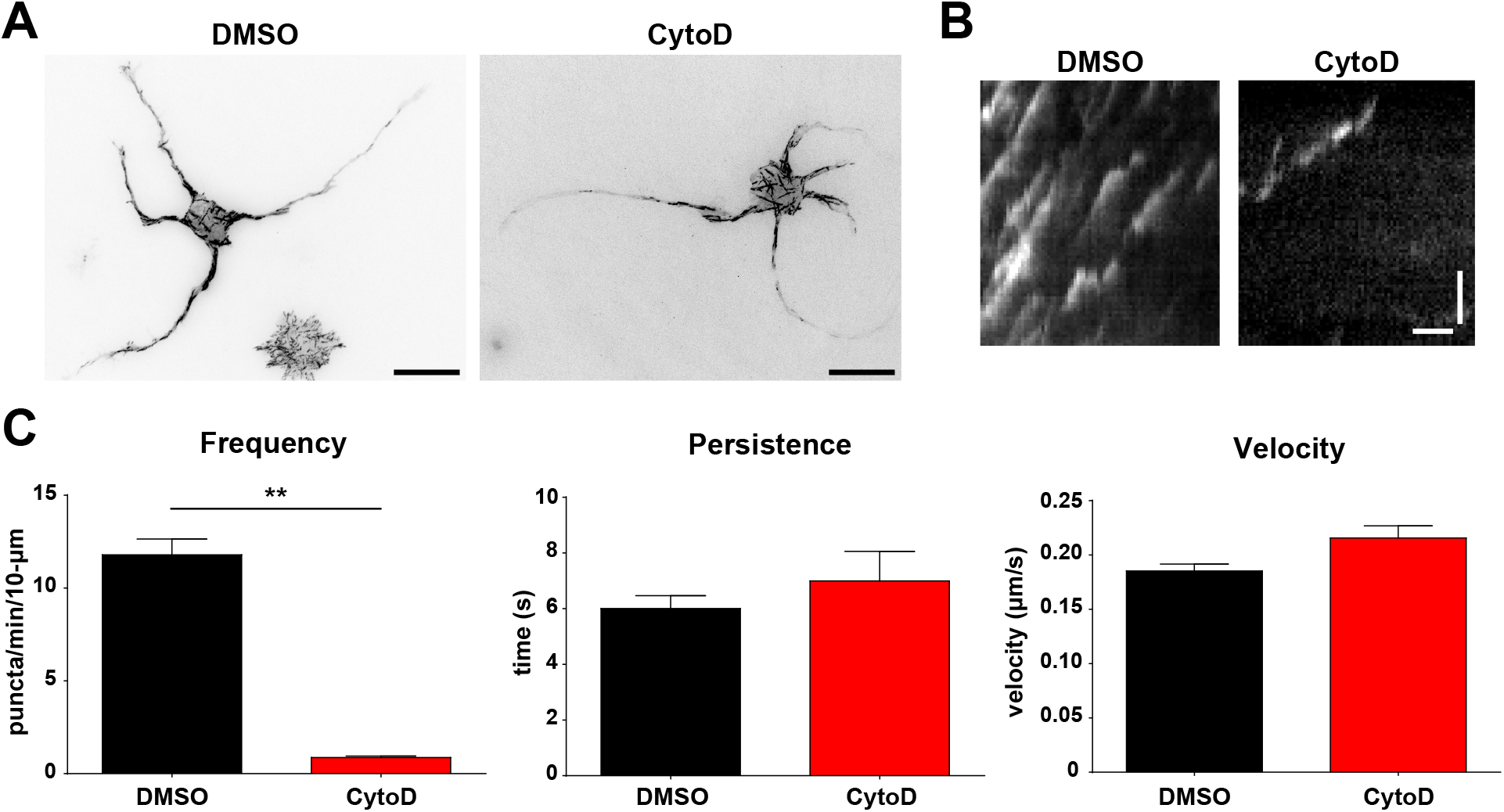
Disrupting actin waves reduces non-centrosomal microtubule nucleation at the neurite tip. (A) Representative maximum projection images (over 2 minute period) of 1DIV EB3-mCherry-expressing hippocampal neurons treated with DMSO (left) or 2.5 μM cytochalasin D (right) for 6 hours. Hippocampal neurons were transfected with plasmids expressing EB3-mCherry and EGFP immediately before plating, incubated for 18 hours, and treated with DMSO or 2.5 μM cytochalasin D for 6 hours before subjected to live cell imaging. The scale bars represent 20 μm. (B) Representative kymographs of EB3-mCherry at the neurite tip in DMSO- (left) or 2.5 μM cytochalasin D-treated (right) neurons. The vertical scale bar in the kymograph represents 2 μm and horizontal scale bar represents 10 seconds. (C) Quantification of EB3-mCherry dynamics at the neurite tip in DMSO- or 2.5 μM cytochalasin D-treated neurons. ** *p*<0.01, two-tailed Student’s *t*-test. Error bars represent SEM from three independent experiments, with more than 45 neurites analyzed in each condition.

## Materials and Methods

### Antibodies and reagents

Anti-RanGTP antibody (AR-12) was a kind gift from Ian Macara (Richards et al., 1995). Mouse anti-β-III-tubulin antibody TUJ1 (801202) and mouse anti-neurofilament monoclonal antibody SMI312 (837904) were from Biolegend (San Diego, CA). Mouse anti-α-tubulin antibody (DM1A) was from Millipore (Billerica, MA). Alexa Fluor-conjugated phalloidin, Alexa Fluor-conjugated secondary antibodies, and MitoTracker Red were from Thermo Fisher Scientific (Waltham, MA).

### Expression plasmids construction

Ran expressing plasmids (pCAG-AcGFP-RanWT, pCAG-AcGFP-RanT24N) were cloned by inserting wild-type Ran or RanT24N obtained from pWT-Ran-V5-His-TOPO or pT24N-Ran-V5-His-TOPO using KpnI and PmeI into KpnI- and SmaI-digested pCAG-AcGFP1-C3 vector. pWT-Ran-V5-His-TOPO and pT24N-Ran-V5-His-TOPO were kindly provided by Richard Cerione (Ly et al., 2010). pCAG-AcGFP-RanQ69L were cloned by inserting RanQ69L obtained from pmCherry-C1-RanQ69L using BamHI and HindIII into pCAG-AcGFP1-C1 digested with the same enzymes. pmCherry-C1-RanQ69L (Addgene plasmid # 30309) was a kind gift from Jay Brenman (Kazgan et al., 2010).

pTriEx-NTOM20-mVenus-LOV2wt, pTriEx-mCherry-ZDK1, and pTriEx-PA-Rac1-I539E (Addgene plasmid # 22026) were kindly provided by Klaus Hahn (Wang et al., 2016). pTriEx-NTOM20-mVenus-LOV2-I539E was constructed by PCR amplifying LOV2-I539E from pTriEx-PA-Rac1-I539E, digested with BamHI/EcoRV, and ligated to BamHI/HindIII digested pTriEx-NTOM20-mVenus-LOV2wt vector. The HindIII cut site of BamHI/HindIII digested pTriEx-NTOM20-mVenus-LOV2wt vector was blunted by Klenow fragment enzyme before ligation. pTriEx-mCherry-ZDK1-RanQ69L and pTriEx-mCherry-ZDK1-RanT24N plasmids were cloned by inserting PCR amplified RanQ69L from pmCherry-C1-RanQ69L and RanT24N from pCAG-AcGFP-RanT24N into HindIII/XhoI digested pTriEx-mCherry-ZDK1 plasmid. A stop codon between ZDK1 and RanQ69L (or RanT24N) was removed by NotI/HindIII digestion, followed by ends blunting using Klenow fragment and self-ligation.

### Neuron culture and transfection

All animal experimental procedures were approved by the Institutional Animal Care and Use Committee (IACUC) and in accordance with the Guide for the Care and Use of Laboratory Animals of National Chiao Tung University. Dissociated hippocampal and cortical neuron cultures were prepared as previously described with slight modification (Chen et al., 2017). Briefly, hippocampi or cortexes from E17.5 mouse embryos were dissected, digested with trypsin-EDTA, and triturated. Dissociated neurons were seeded onto poly-L-lysine-coated coverslip (2.5×10^3^ cells/cm^2^ for low-density cultures and 3×10^4^ cells/cm^2^ for regular culture). Plasmids were introduced into neurons using Nucleofector II (Lonza, Basel, Switzerland) immediately before seeding or using lipofectamine 2000 (Thermo Fisher Scientific, Waltham, MA) at indicated days *in vitro*. Lipofectamine transfected cells were incubated for 4 hours and the medium containing transfection mixture was replaced with cortical neuron-conditioned neurobasal medium (low-density cultures) or fresh neurobasal medium plus B27 supplement (regular culture).

### HeLa cell synchronization and transfection

7×10^4^ HeLa cells were seeded per well in a 24-well plate. 6~8 hours after seeding, the growth medium was replaced with 2 mM thymidine-containing growth medium and incubated for 16 hours. After this first thymidine block, cells were released and allowed to progress through cell cycle for 4 hours. At this point, lipofectamine 2000-based transfection was performed. 4 hours post-transfection, the growth medium was again replaced with 2 mM thymidine-containing growth medium and incubated for another 16 hours. After the second thymidine block, HeLa cells were released and incubated for 10 hours before being fixed for subsequent analyses.

### Indirect immunofluorescence staining

Cells were fixed with 3.7% formaldehyde for 15 minutes at 37 °C and then washed three times with PBS. Fixed cells were permeabilized with 0.25% triton X-100 in PBS for 5 minutes at room temperature and washed with PBS three times. Cells were then blocked with 10% BSA in PBS for 30 minutes at 37 °C. Coverslips with cells were then incubated for 1 hour at 37 °C with different primary antibodies: anti-α-tubulin (1:1000), anti-β-III-tubulin (1:4000), anti-RanGTP (1:100), and anti-neurofilament (1:1000). After primary antibody incubation, cells were incubated with Alex Fluor-conjugated secondary antibodies (1:1000). All antibodies were diluted in 2% BSA in PBS. Coverslips with cells were washed with PBS three times and mounted with Fluoromount onto glass slides.

### Image acquisition

Immunofluorescence stained HeLa cells were acquired on an Olympus IX-71 inverted microscope equipped with a 40× 0.95 N.A. Plan Apochromat objective lens, a CoolLED epi-fluorescence light source, a Hamamatsu ORCA-R2 camera, and MetaMorph software 7.6.5.0. Immunofluorescence stained neurons were acquired on a Nikon Eclipse-Ti inverted microscope equipped with a 60× 1.49 N.A. Plan Apochromat objective lens, an Intensilight epi-fluorescence light source, a Photometrics CoolSNAP HQ2 camera, and Nikon NIS-Elements software 4.13.05.

Live cell imaging was performed on a Nikon Eclipse-Ti inverted microscope equipped with a 60× 1.49 N.A. Plan Apochromat objective lens, a Photometrics CoolSNAP HQ2 camera, a built-in Perfect Focus system, a Tokai Hit TIZHB live cell chamber, and Nikon NIS-Elements software 4.13.05. For quantifying MT dynamics, EB3-mCherry was excited using a TIRF illuminator and a 561 nm DPSS laser; images were acquired every 500 milliseconds over 1- or 2-minute period. For examining actin waves and the migration of Ran mutants along the neurite, AcGFP-Ran molecules were excited using an Intensilight epi-fluorescence light source; both DIC and fluorescent images were acquired every 3 minutes over a 4-hour period.

### Photoactivation of RanTRAP

Photoactivation experiments were performed on the same microscope as live cell imaging experiments. A 60× 1.49 N.A. Plan Apochromat objective lens and an Intensilight epi-fluorescence illuminator were used for photoactivation. The field diaphragm was closed down to the minimal size (~40 μm in diameter) during the photoactivation period. A Nikon stock FITC filter set and the 1/32 ND filter were used to condition the photoactivation light. A 24-pulse regime (24 pulses of 80 milliseconds irradiation time with 5 seconds interval between pulses) was performed using NIS-Element software 4.13.05. All photoactivation regions were selected more than 15 μm away from the soma or the tip of the neurite.

### Image analysis

For analyzing the distribution of RanGTP signals along the neurite (linescan analyses), only neuronal protrusions that are β-III-tubulin-positive, and are longer than the diameter of its soma were considered as neurites. In addition, only neurites not intersecting others were included in the analysis. The analysis was performed manually by tracing neurites from the soma edge to the neurite tip (neurites without growth cone) or to the wrist of the growth cone (neurites with growth cone) with a segment line 0.55 μm in width in the RanGTP signal channel using ImageJ 1.49v.

For EB3-mCherry comets analysis, NIS-Elements software 4.13.05 was used to generate the kymograph for the EB3-mCherry channel. A window 40 μm (for photoactivation experiments) or 10 μm (for all other experiments) in length and 0.77 μm in width was used to generate the kymograph. Only EB3-mCherry movements that could be followed clearly for equal or more than 4 frames (1.5 seconds) were defined as an event. The emanating frequency of EB3-mCherry was quantified from the kymograph by manually counting the number of EB3-mCherry events per minute. The velocity and persistence time of EB3-mCherry were quantified from the kymograph by drawing a line along an EB3-mCherry event.

For Ran mutant and actin wave co-migration analysis, only neurites with actin waves were included in the analysis. The position of the actin wave along the neurite shaft was manually determined from the DIC images. The analysis was performed by first manually tracing neurites from the edge of the soma to the neurite tip (for neurites without growth cone) or to the wrist of the growth cone (for neurites with growth cone) with a segment line 0.55 μm in width. This manually traced line was then used to perform linescan analysis in the AcGFP-Ran channel or the DIC channel using ImageJ 1.49v. The signal intensity of AcGFP-Ran was normalized so that the highest intensity along the manually traced segment is set to 1 and lowest intensity to 0.

For quantifying AcGFP-Ran intensity change over time, a circular ROI (4-pixel in diameter) was created at the center of an actin wave-containing neurite using ImageJ 1.49v. The intensity of AcGFP in this ROI was measured every 3 minutes from 15 minutes before the arrival of the actin waves (−15 min) to 15 minutes after the actin wave has passed (15 min). The intensity of AcGFP was normalized to −15 minutes time point.

### Statistical analysis

All statistical analyses were performed using GraphPad Prism 7 with the indicated statistical methods.

## Discussion

While recent works have started to uncover the cellular components crucial for nucleating ncMTs in neurons, no regulatory mechanism has yet to be discovered. Here we show that the small GTPase Ran plays an important role in controlling ncMT nucleation along the neurite. We first show that expressing the RanGTP- or RanGDP-mimic mutant can specifically increase or decrease the RanGTP level at the tip of the neurite, this in turn enhances or reduces ncMT nucleation at the neurite tip. To demonstrate that Ran can regulate ncMT nucleate in regions other than the neurite tip, we constructed an optogenetic tool called RanTRAP that allows the release of the RanGTP-mimic mutant at the photoactivation site. By photoactivating RanTRAP along the neurite shaft where endogenous RanGTP level is low, we demonstrate that Ran GTPase can indeed control ncMT nucleation in neurons. In addition, we identified a novel transport mechanism for RanGTP in neuronal cytoplasm. First, it was observed that the endogenous RanGTP colocalizes with actin waves in fixed neurons. Second, disrupting the actin wave propagation with cytochalasin D essentially eliminates RanGTP enrichment at the neurite tip. Finally, the co-migration of the AcGFP-tagged RanGTP-mimic mutant and the actin wave can be observed in live neurons. These observations demonstrate that the actin wave is a major contributor to the enrichment of RanGTP at neurite tips. Consistently with the idea that RanGTP promotes ncMT nucleation and it is transported by actin waves to the neurite tip, cytochalasin D treatment which abolishes actin waves also causes a significant reduction of ncMT nucleation at the neurite tip.

Given that γ-tubulin, augmin, and TPX2 interact with each other and that MT nucleation from the sides of existing MTs requires all three proteins in *Xenopus* egg extract (Petry et al., 2013), it is tempting to speculate that RanGTP promotes ncMT nucleation by releasing TPX2 from the inhibitory importin heterodimers which in turn activating the γ-tubulin-augmin-TPX2 complex at the neurite tip. In support of this hypothesis, TPX2 and augmin complex subunits have been observed to localize along neurites (Chen et al., 2017; Cunha-Ferreira et al., 2018). In addition, both importin-α and importin-β molecules have been observed to present throughout the cytoplasm in cultured neurons (Chen et al., 2017; Hanz et al., 2003). Furthermore, the Ran-importin-β complexes have been detected in the soma and the neurite tips of hippocampal neurons (Chen et al., 2017). It will be interesting to test whether the γ-tubulin-augmin-TPX2 complex exists along the neurite and whether Ran also regulates augmin-mediated ncMT nucleation.

The observation that locally releasing RanGTP along the neurite shaft leads to ncMT nucleation at the releasing site suggests neurons may utilize Ran to dynamically reorganize their MT network. It is currently unclear if and how neurons can control the localization of Ran in different regions. One possibility is the utilization of RanGTP-anchoring protein(s) to trap RanGTP at specific regions of the neuron. For example, RanBP9 is a RanGTP-binding protein that has been shown to causes ectopic MT nucleation when overexpressed (Nakamura et al., 1998). In addition, RanBP9 has been observed to localize along the neurite (Lakshmana et al., 2012). Considering Ran depletion leads to morphological changes in neurons (Mencarelli et al., 2018; Sepp et al., 2008), it is likely that locally elevating RanGTP level can also alter the morphology of the neuron. It will be interesting to test this hypothesis by long-term RanQ69L-TRAP photoactivation along the neurite shaft to see if this can lead to the formation of a collateral branch.

The discovery that RanGTP is transported by actin waves towards the neurite tip provides a novel mechanism for cells to position the cytosolic small GTPase. While the effect of Ran GTPase on positioning actin cytoskeleton has previously been observed in *Xenopus* egg (Deng et al., 2007; Yi et al., 2011), the effect of actin cytoskeleton on positioning Ran GTPase has never been described. It is interesting to point out that the increase of MT nucleation and MT density can be detected immediately behind the actin wave (Winans et al., 2016), it is hypothesized that actin waves make the neurite wider to create space for more MTs to form. Perhaps the increase of MT nucleation and density can also be attributed to the effect of RanGTP moving with the actin wave, causing ncMT formation in its wake. In order for RanGTP to be transported by the actin wave, specific adapter protein(s) that localizes to the actin wave and interact with RanGTP must be present. As of now, the identity of the adapter protein(s) is unknown. One potential candidate of this adapter is the protein ezrin, an actin-binding protein that is concentrated within actin waves (Ruthel and Banker, 1998). Ezrin was later discovered to interact with the cytoplasmic domain of L1CAM (Dickson et al., 2002), a neural cell adhesion molecule that interacts with the Ran-binding protein RanBP9 to regulate neurite outgrowth (Cheng et al., 2005; Woo et al., 2012). RanBP9 was found to selectively bind to RanGTP in a yeast 2-hybrid screen (Nakamura et al., 1998). Collectively, these data suggest an ezrin-L1CAM-RanBP9 complex to act as the adapter to transport RanGTP within actin waves. In addition to adapter proteins in the actin wave, we believe specific RanGTP-anchoring protein(s) exist at the neurite tip in order for RanGTP to be enriched at this location. This is because the frequency of actin wave generation (1~2 waves per hour) is much too low to maintain an elevated pool of cytosolic RanGTP at the neurite tip (Winans et al., 2016). Second, the frequency of actin waves decreases after neurons have been cultured for 3~4 days *in vitro* (Flynn et al., 2009; Ruthel and Banker, 1999), yet RanGTP remains enriched at the neurite tip in later neurons (data not shown). One interesting candidate of this anchoring protein is RanGAP1, which has been observed to localize to the tips of growing axons in DRG neurons (Yudin et al., 2008). Given that importin-β has been shown to inhibit the RanGAP1-stimulated hydrolysis of RanGTP (Floer and Blobel, 1996) and the observation of importin-β-Ran complex in the neuronal cytoplasm (Chen et al., 2017), it is possible that the RanGAP1 can anchor RanGTP at the neurite tip without catalyzing the hydrolysis of GTP. Using RanGAP1 as the anchoring protein also provides a convenient inactivation mechanism once the need for a RanGTP-enriched neurite tip is no longer required.

## Acknowledgments

We thank Prof. Klaus Hahn for providing the pTriEx-NTOM20-mVenus-LOV2wt and pTriEx-mCherry-ZDK1 plasmids. We are grateful to Prof. Richard Cerione for providing the pWT-Ran-V5-His-TOPO and pT24N-Ran-V5-His-TOPO plasmids. We also would like to thank Prof. Oliver Gruss and Prof. Ian Macara for providing the anti-TPX2 and anti-RanGTP antibodies, respectively. This work was supported by grants from Taiwan Ministry of Science and Technology (MOST 105-2320-B-009-005-MY3) and the “Center for Intelligent Drug Systems and Smart Bio-devices (IDS2B)” from The Featured Areas Research Center Program within the framework of the Higher Education Sprout Project by the Ministry of Education (MOE) in Taiwan.

