## Supplementary Data for "Ran GTPase regulates non-centrosomal microtubule nucleation and is transported by actin waves towards the neurite tip"

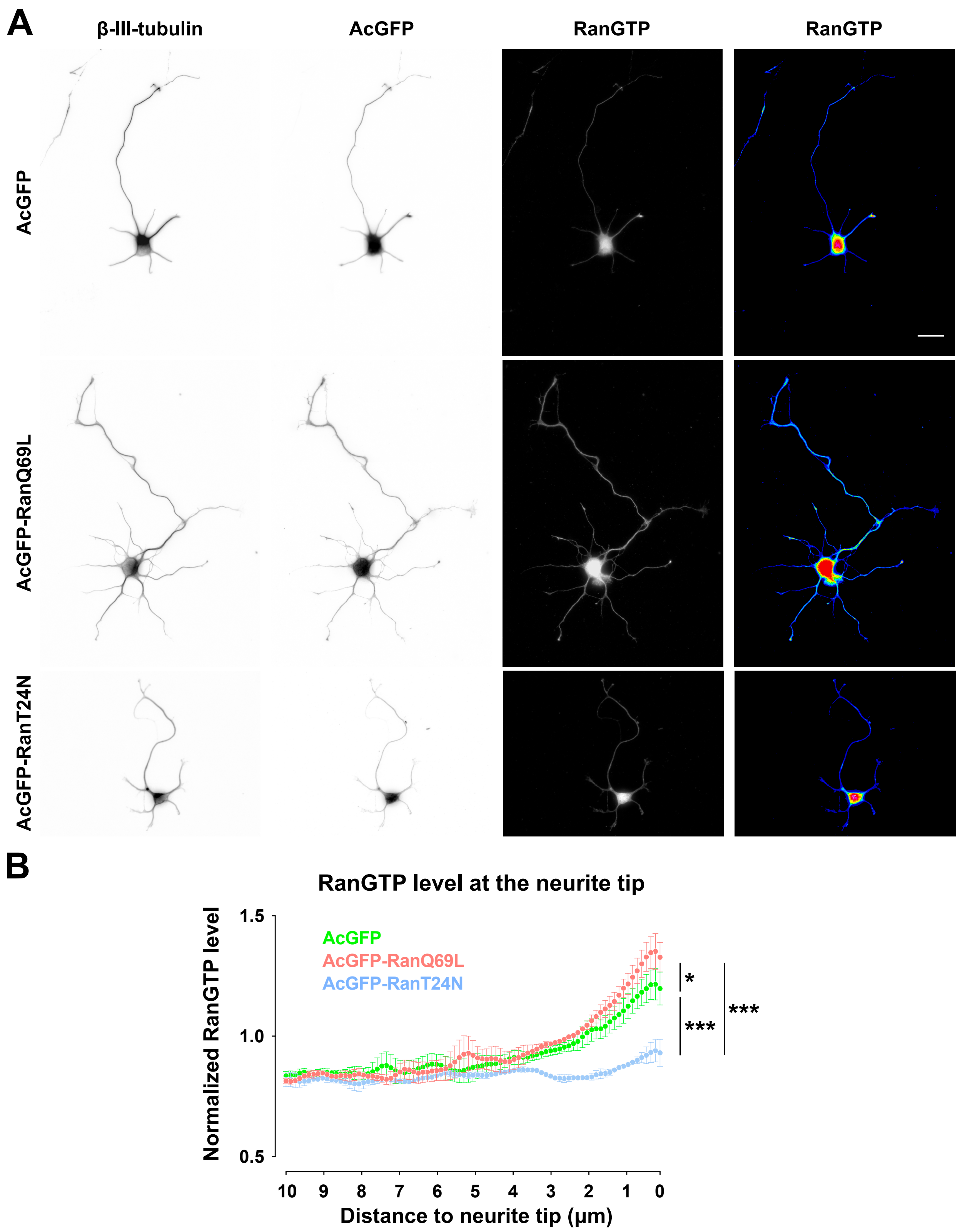
A
β-III-tubulin
AcGFP
RanGTP
RanGTP
AcGFP
AcGFP-RanQ69L
AcGFP-RanT24N
B
RanGTP level at the neurite tip
Normalized RanGTP level
1.5
1.0
0.5
AcGFP
AcGFP-RanQ69L
AcGFP-RanT24N
*
***
***
10
9
8
7
6
5
4
3
2
1
0
Distance to neurite tip (μm)

**Figure S1. Ran mutants influence the level of RanGTP at the neurite tip.**

(A) Representative images of 2DIV low-density dissociated hippocampal neurons expressing AcGFP, AcGFP-RanQ69L, or AcGFP-RanT24N. β-III-tubulin, AcGFP, RanGTP, and pseudo-colored RanGTP channels are shown. All images have the same scale and the scale bar represents 20 μm. (B) Normalized RanGTP level linescans at the neurite tip (AcGFP is shown in green, AcGFP-RanQ69L in red, and AcGFP-RanT24N in blue). RanGTP intensity 0~10 μm from the neurite tip was normalized to the average RanGTP intensity along the entire neurite. The dots and error bars indicate mean and SEM from 3 independent experiments. At least 119 neurites from 3 independent experiments from each condition were analyzed. *** $p<0.001$, * $p<0.05$, two-way ANOVA followed by Tukey post-hoc test (RanGTP level 0~1 μm from the tip of the neurite).

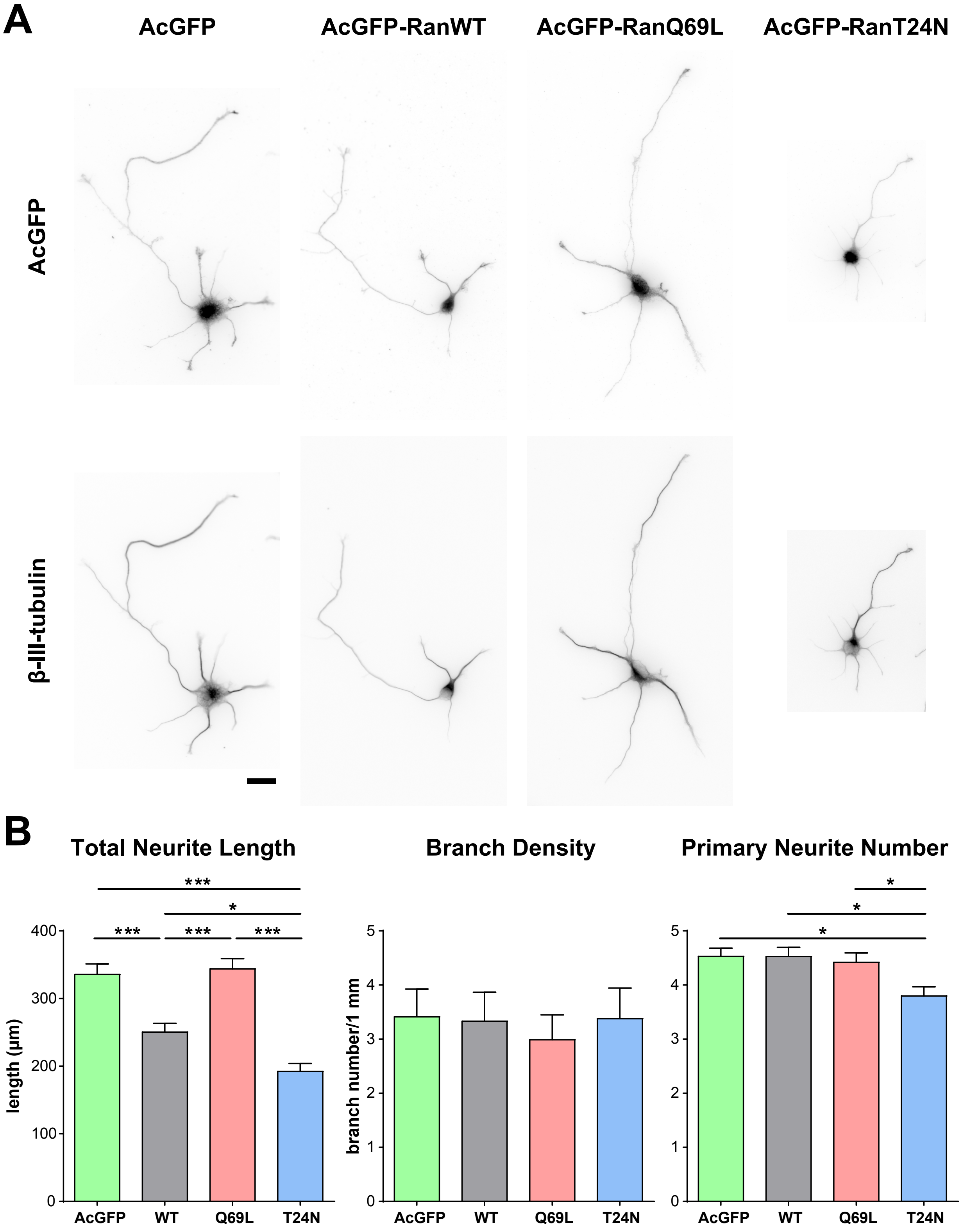
A
AcGFP
AcGFP-RanWT
AcGFP-RanQ69L
AcGFP-RanT24N
AcGFP
β-III-tubulin
B
Total Neurite Length
Branch Density
Primary Neurite Number
length (μm)
branch number/1 mm
AcGFP
WT
Q69L
T24N
***
*
0
100
200
300
400
0
1
2
3
4
5

**Figure S2. Ran mutants affect neuronal morphogenesis.**

(A) Representative images of 2DIV low-density dissociated hippocampal neurons expressing AcGFP, AcGFP-RanWT, AcGFP-RanQ69L, or AcGFP-RanT24N. AcGFP (top) and β-III tubulin (bottom) channels are shown. All images have the same scale and the scale bar represents 20 μm. (B) Quantification of total neurite length (left), branch density (middle), and primary neurite number (right) in Ran expressing neurons. * $p<0.05$, *** $p<0.001$, one-way ANOVA followed by Tukey's post-hoc test. Error bars represent SEM from 3 independent experiments.

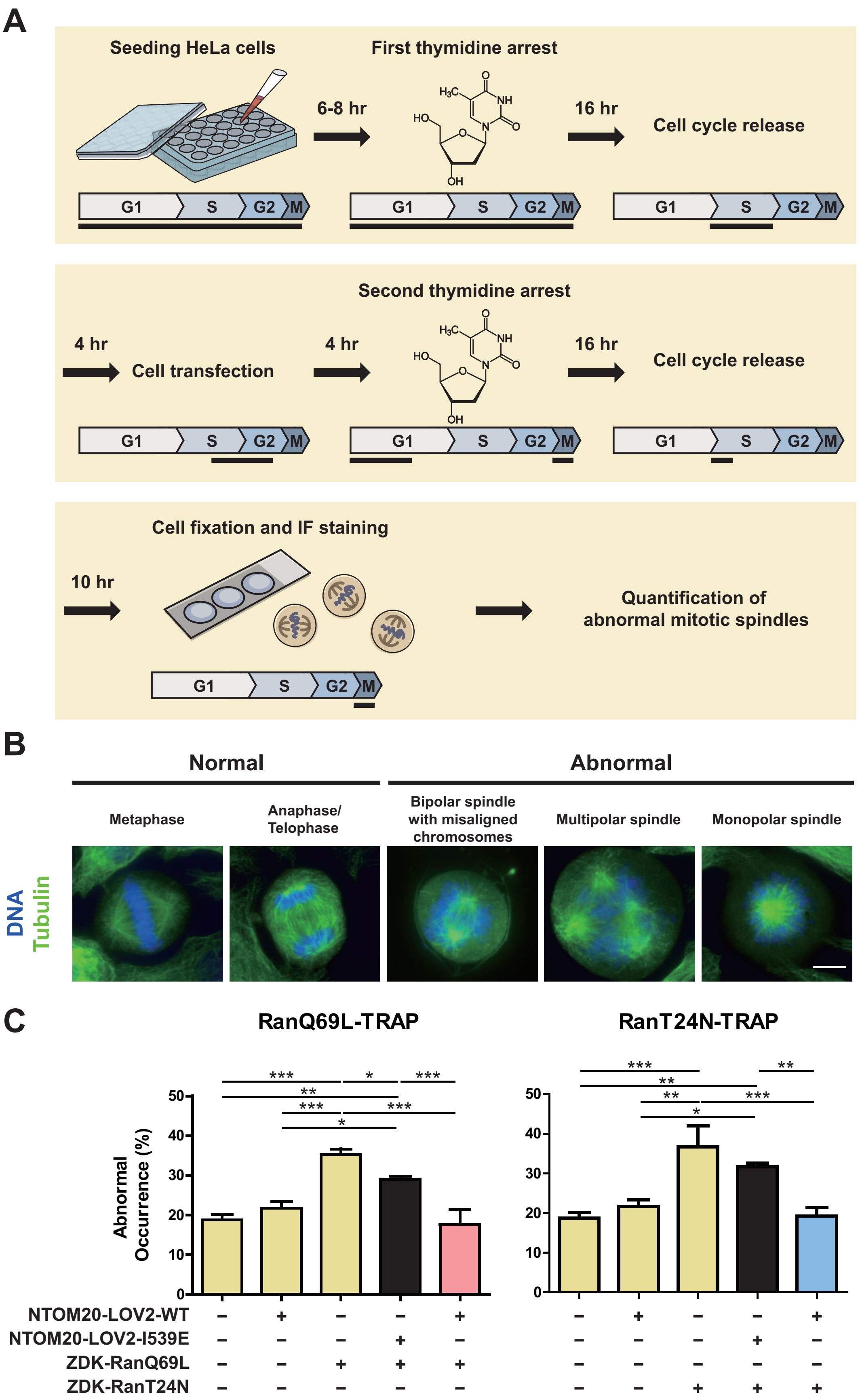

A
Seeding HeLa cells
First thymidine arrest
6-8 hr
16 hr
Cell cycle release
G1
S
G2
M
Second thymidine arrest
4 hr
Cell transfection
4 hr
16 hr
Cell cycle release
Cell fixation and IF staining
10 hr
Quantification of abnormal mitotic spindles
B
Normal
Abnormal
Metaphase
Anaphase/ Telophase
Bipolar spindle with misaligned chromosomes
Multipolar spindle
Monopolar spindle
DNA
Tubulin
C
RanQ69L-TRAP
RanT24N-TRAP
Abnormal Occurrence (%)
NTOM20-LOV2-WT
NTOM20-LOV2-I539E
ZDK-RanQ69L
ZDK-RanT24N

**Figure S3. Functional validation of RanTRAP system in HeLa cells.**

(A) Schematic procedure of double thymidine arrest to synchronize HeLa cells. The black horizontal bars indicate where in the cell cycle most cells are. (B) Representative images of synchronized mitotic HeLa cells with normal or abnormal spindles. Cells were stained with α-tubulin antibody (DM1A) and DAPI. The spindle morphology was categorized using the following criteria. A mitotic spindle containing two spindle poles and aligned chromosomes was categorized as "metaphase" spindle. A mitotic spindle containing two spindle poles and segregated chromosomes was categorized as "anaphase/telophase" spindle. A mitotic spindle containing two spindle poles and misaligned chromosomes was categorized as "bipolar spindle with misaligned chromosomes". A mitotic spindle containing more than two spindle poles was categorized as "multipolar spindle". A mitotic spindle containing only one spindle pole and radially arranged chromosomes was categorized as "monopolar spindle". The scale bar represents 10 μm. (C) Quantification of abnormal spindle occurrence. More than 50 mitotic HeLa cells were analyzed per condition per repeat. * $p<0.05$, ** $p<0.01$, *** $p<0.001$, one-way ANOVA followed by Tukey's multiple comparison test. Error bars represent SEM from 3 independent experiments.

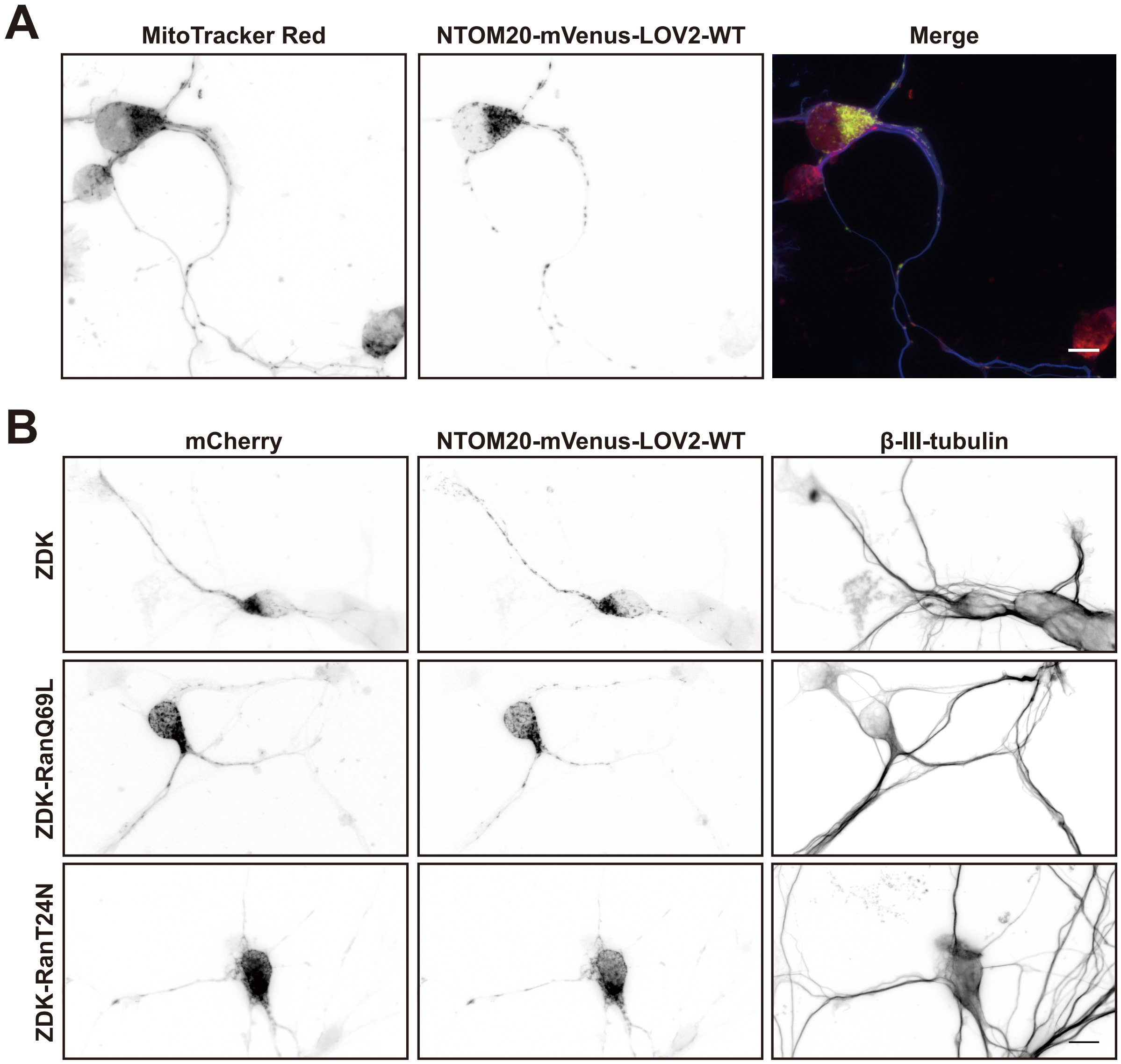


**Figure S4. The RanTRAP system targets Ran to mitochondria along the neurite.**

(A) Representative images of a mouse cortical neurons transfected with plasmid expressing NTOM20-mVenus-LOV2-WT (green in merged) at 2DIV, fixed and stained with MitoTracker Red (red in merged) and β-III-tubulin antibody (blue in merged) at 4DIV. (B) Representative images of mouse cortical neurons transfected with plasmids expressing NTOM20-mVenus-LOV2-WT and mCherry-ZDK (or mCherry-ZDK-RanQ69L or mCherry-RanT24N) at 2DIV, fixed and stained with the β-III-tubulin antibody at 4DIV. All scale bars represent 10 μm.

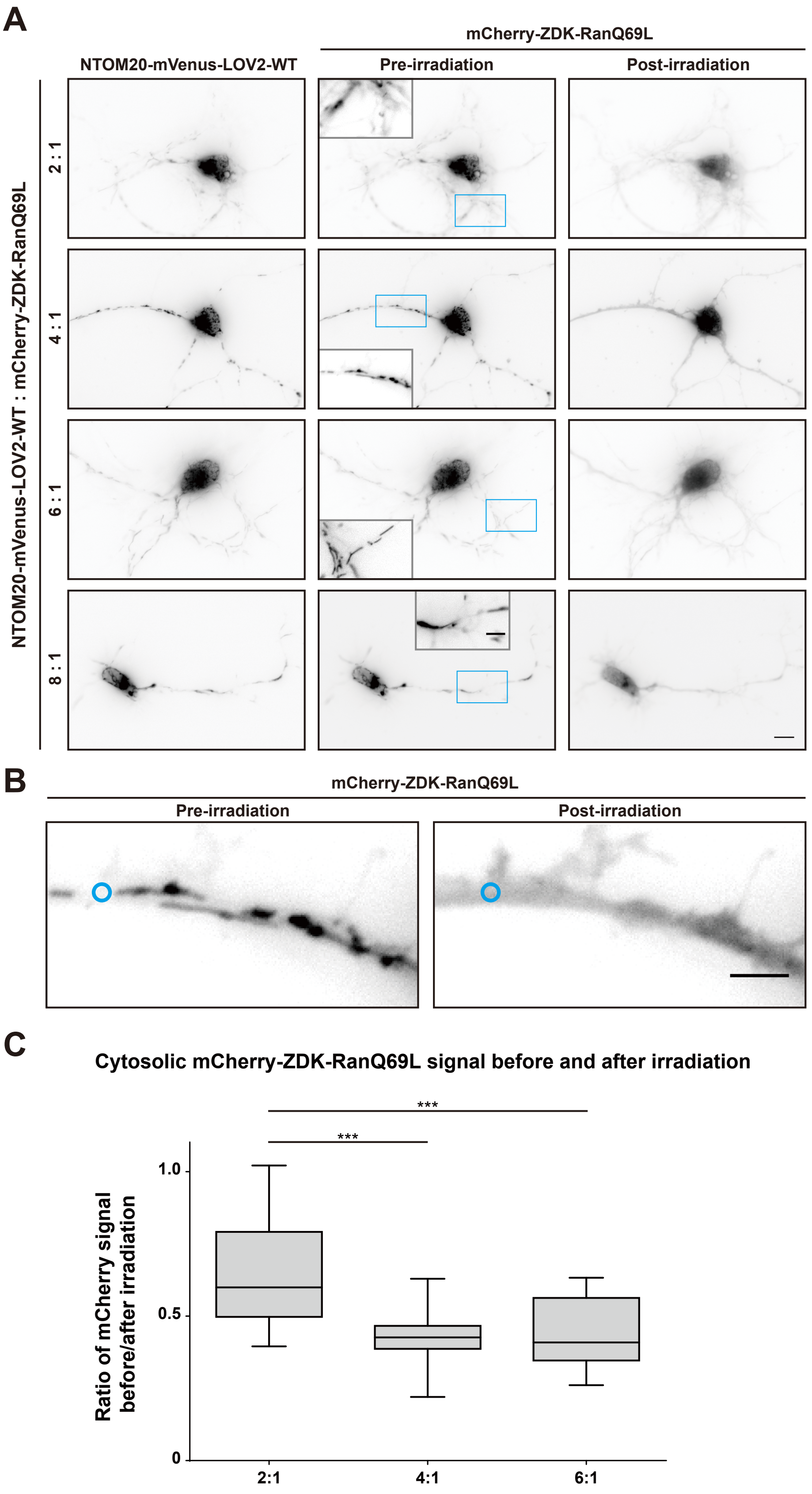

A
NTOM20-mVenus-LOV2-WT
mCherry-ZDK-RanQ69L
Pre-irradiation
Post-irradiation
NTOM20-mVenus-LOV2-WT : mCherry-ZDK-RanQ69L
2 : 1
4 : 1
6 : 1
8 : 1
B
mCherry-ZDK-RanQ69L
Pre-irradiation
Post-irradiation
C
Cytosolic mCherry-ZDK-RanQ69L signal before and after irradiation
***
***
Ratio of mCherry signal before/after irradiation
1.0
0.5
0
2:1
4:1
6:1

**Figure S5. Optimizing the molar ratio of RanTRAP plasmids in neurons.**

(A) Representative live cell images of 4DIV mouse cortical neurons co-expressing NTOM20-mVenus-LOV2-WT and mCherry-ZDK-RanQ69L. Neurons were transfected with various molar ratios of plasmid expressing NTOM20-mVenus-LOV2-WT and mCherry-ZDK -RanQ69L (LOV2:ZDK = 2:1, 4:1, 6:1, 8:1) at 2DIV, incubated for 2 days before subjected to live cell imaging. All images have the same scale and the scale bar represents 10 μm. The image insets show the magnified and contrast enhanced view of the area enclosed by the blue box, and the scale bar in the inset represents 5 μm. (B) Representative images showing the ROI (blue circle) where cytosolic mCherry-ZDK-RanQ69L level was quantified. Both live cell images show 4 DIV mouse cortical neurons expressing NTOM20-mVenus-LOV2-WT and mCherry-ZDK-RanQ69L for 2 days. Blue circular ROIs (1.1 μm in diameter) indicate where cytosolic mCherry-ZDK-RanQ69L signal was quantified from. The scale bar represents 4 μm. (C) Quantification of cytosolic mCherry-ZDK-RanQ69L level before and after photoactivation in selected ROI along the neurite in various molar ratio conditions. In the molar ratio 8:1 condition, most of the transfected neurons are unhealthy. As a result, this condition was excluded from the quantification. 13, 30, and 19 neurites were quantified in 2:1, 4:1, and 6:1 conditions, respectively. Box plot shows median, first and third quartile, and whiskers extend to the entire range of data. *** $p<0.001$, one-way ANOVA followed by Tukey's multiple comparison test. Since no statistically significant difference can be detected between the 4:1 and 6:1 condition, the molar ratio (4:1) that produced the higher signal of mCherry-ZDK-RanQ96L was chosen for subsequent experiments.

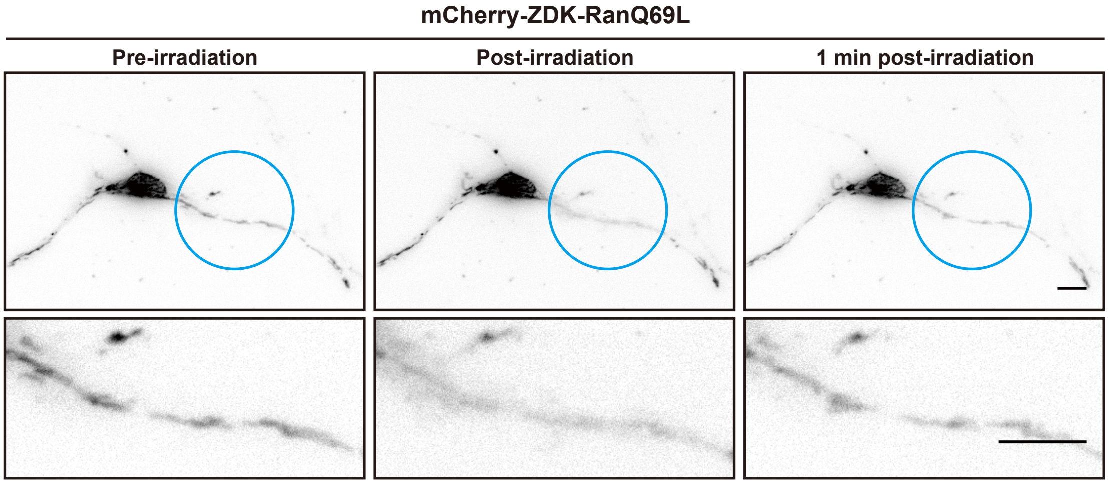


**Figure S6. Local release of RanGTP-mimic mutant can be achieved in neurons.**
Mouse hippocampal neurons were co-transfected with plasmids expressing NTOM20- mVenus-LOV2-WT and mCherry-ZDK-RanQ69L at 3DIV, incubated for 1 days before subjected to live cell imaging. The blue circle indicated the region of photoactivation. Images on the bottom row show magnified images from the photoactivated region. The mCherry-ZDK-RanQ69L signal before (left), immediately after photoactivation (center), and 1 minute after photoactivation (right) are shown. Note that mCherry-ZDK-RanQ69L only dissociated from the mitochondria inside the photoactivated region. All scale bars represent 10 µm.
